# Temporal Gatekeeping Role of Lmx1 during chordate neural tube morphogenesis

**DOI:** 10.64898/2026.03.04.709676

**Authors:** Josefina Pérez-Benítez, Michael S. Levine, Laurence A. Lemaire

**Author notes:** Corresponding author: L.A.L.

## Abstract

Neural tube closure is a critical developmental process, essential to the proper formation of the vertebrate nervous system. This process starts with the invagination of neural plate cells. Its borders then converge, leading to the closure of the neural tube, propagating like a zipper. Afterwards, cell intercalation and proliferation allow the tube to elongate. Neural tube closure involves thousands of cells in vertebrates. However, the closest invertebrates to vertebrates, the tunicates, such as *Ciona*, close a hollow dorsal neural tube with fewer than 20 neural cells. This minimal model makes it easier to study the mechanisms of this intricated process. In *Ciona*, the transcription factor *Lmx1* is expressed in the most dorsal cells of the developing neural tube, like its vertebrate orthologs. In vertebrates, Lmx1 paralogs are involved in neural tube patterning. However, no function related to morphogenesis has been uncovered. Here, we explore *Ciona* Lmx1 roles during neural tube closure. *Lmx1* Knockdown leads to slight but significant defects in neural tube closure. The overexpression of a repressive Lmx1 variant prevents the proper intercalation of the dorsal neural tube cells, impeding the anterior progression of the zipper. Furthermore, studies of *Lmx1* regulatory sequences indicate that Pax3/7, ZicL, and Nodal signaling may directly regulate its transcription. These transcription factors are present at the vertebrate neural plate border, suggesting that *Lmx1* regulation is conserved across chordates. It raises the possibility of an unrecognized role for Lmx1 during vertebrate neural tube morphogenesis.

**Graphical abstract:** 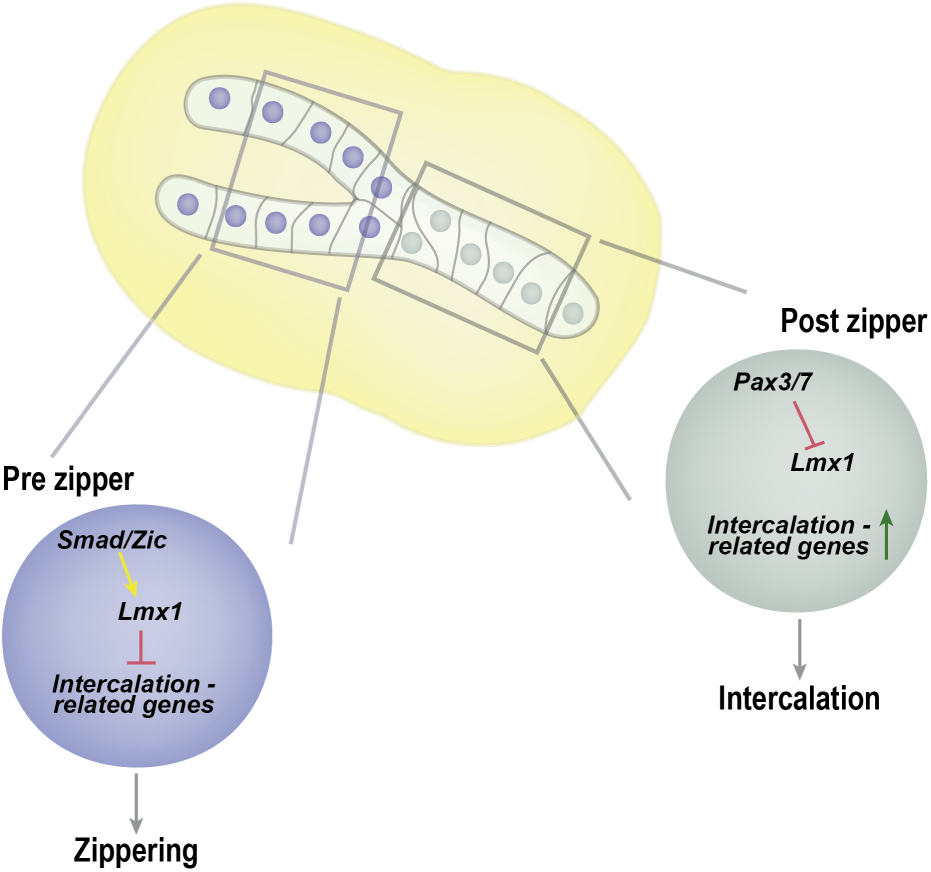

## 1. Introduction

The chordate central nervous system arises from a tube formed during neurulation. At that time, the neural plate bends at the midline, and its edges elevate, forming the neural folds. The folds then converge toward the midline, pushed by internal apical constriction at the hinge points and external convergence extension forces from the surface ectoderm. The folds progressively fuse to form a hollow tube. In mammals, closure occurs at multiple initiation points and progresses in a zipper-like fashion along the antero-posterior axis of the embryo. This tube subsequently elongates, driven by convergence extension and cell proliferation (Nikolopoulou et al., 2017). Defects in neural tube closure are the second most common congenital malformation in humans, affecting approximately 1 in 1000 individuals worldwide with substantial geographical disparities (Zaganjor et al., 2016). Neural tube closure defects are classified based on the location of the failure. It manifests as anencephaly, when the tube remains open at the cranial level, or spina bifida, when the defect occurs at the level of the spinal cord. These conditions are multifactorial, resulting from a confluence of genetic and environmental factors, impeding the elucidation of their pathogenicity (Nikolopoulou et al., 2017).

This complex multi-step process is controlled by numerous molecular actors. BMP and Shh signaling pathways control the formation of hinge points, allowing the neural groove to form and the neural folds to converge (Ybot-Gonzalez et al., 2002; Ybot-Gonzalez et al., 2007). Impairments in Wnt and the planar cell polarity signaling pathway disrupt convergence extension and also cause neural tube defects (Ossipova et al., 2015; Wallingford and Harland, 2002). Finally, transcription factors present in the neural folds, such as *Cdx*, *Zic2*, and *Pax3*, or in the surface ectoderm, like the *Grhl* transcription factors, regulate different aspects of the closure, from controlling specific signaling pathways and hinge point formation to cell cycle and adhesion regulation (Escuin et al., 2023; Pyrgaki et al., 2011; Savory et al., 2011; Sudiwala et al., 2019). While the intricacies of these pathways and molecular mechanisms are increasingly well-characterized, their interactions and temporal coordination only start to be uncovered (Nagaoka et al., 2023; Nychyk et al., 2022; Palmer et al., 2021).

Given that *Ciona* belongs to the closest living invertebrate group to the vertebrates (Bourlat et al., 2006; Delsuc et al., 2006; Delsuc et al., 2008), its embryo provides a powerful model tube to study the spatial and temporal coordination behind neural tube morphogenesis. The central nervous system of *Ciona* larvae develops from a hollow dorsal tube, whose formation evokes vertebrate neurulation on a minimalistic scale. The neural plate is only 6 to 8 cells wide. The cells invaginate, and the neural plate borders converge toward the midline, where they fuse. The closure point initiates on the posterior side of the embryo, starting at the blastopore, and progresses anteriorly like a zipper (Nicol and Meinertzhagen, 1988). On the other side of the tube, the neuropore closes by apical constriction (Veeman et al., 2010).

The zipper only moves through 9 to 10 cells on both sides of the groove. These cells always have the same developmental history owing to the embryo’s stereotypical lineages. On each side of the neural plate, the 5 posterior cells originate from the b-blastomeres at the 8-cell stage (Cole and Meinertzhagen, 2004; Conklin, 1905; Nicol and Meinertzhagen, 1988). On the molecular level, the zipper progresses via cell junction shortening through actomyosin contraction and rearrangement of cadherins (Hashimoto and Munro, 2019; Hashimoto et al., 2015). Once the zipper occurs, the newly positioned dorsal cells rapidly intercalate before undergoing mitosis (Cole and Meinertzhagen, 2004; Nicol and Meinertzhagen, 1988). The transcriptional regulation underlying these later stages remains elusive.

*Ciona* dorsal neural tube expresses factors conserved with vertebrates, including *Pax3/7, Cdx, Msx,* and *Lmx1* (Beck et al., 1995; Chizhikov and Millen, 2004a; Goulding et al., 1991; Graham et al., 1993; Imai et al., 2004; Imai et al., 2006; Jostes et al., 1990; Meyer and Gruss, 1993; Millen et al., 2004). *Lmx1* is the orthologs of vertebrate *Lmx1a* and *Lmx1b*. These factors act redundantly and are necessary for the neural tube roof patterning. However, their absence does not affect the tube’s closure (Chizhikov and Millen, 2004a, b; Mishima et al., 2009). At later stages, *Lmx1* factors regulate the proliferation-differentiation balance of dopaminergic neuron precursors in the mouse midbrain (Yan et al., 2011). These data show that Lmx1 can coordinate differentiation and cell cycle. However, no role linking them to morphogenesis has been uncovered.

We recently characterized the *Ciona Lmx1* expression pattern at the single-cell resolution. In *Ciona*, *Lmx1* is rapidly downregulated after zippering, suggesting it may be involved in tube closure rather than later patterning. Moreover, overexpression experiments show that it could promote proliferation. Nevertheless, the future dorsal cells expressing *Lmx1* do not divide before the zipper happens (Ostlund-Sholars et al., 2025). Therefore, in this study, we investigated the role of Lmx1 in the temporal coordination of *Ciona* neural tube morphogenesis. We present evidence that *Lmx1* is precisely regulated by the conserved factors Pax3/7 and Zicl, as well as Nodal signaling. Building on our previous study of *Lmx1* involvement in neural tube morphogenesis (Ostlund-Sholars et al., 2025), we show that *Lmx1* is necessary for robust neural tube closure. Strikingly, neural tube closure defects were associated with the impairment of cellular intercalation, which normally occurs after the closure when *Lmx1* is no longer expressed. These results reveal that *Lmx1* might act as a temporal gatekeeper, preventing the early expression of genes involved in later steps of neural tube morphogenesis.

## 2. Results and Discussion

### 2.1. Transcriptional control of Lmx1

In *Ciona*, *Lmx1* expression is tightly regulated in the future dorsal neural tube during closure. *Lmx1* is expressed before the zipper occurs. However, it is rapidly downregulated after passing of the zipper (Ishida and Satou, 2024; Ostlund-Sholars et al., 2025). To understand *Lmx1* transcriptional regulation, we further analyzed the upstream sequences of *Lmx1* that recapitulate its expression (Lemaire et al., 2021). A minimal enhancer of 300bp active in the dorsal neural cells was identified by performing 5’ truncation of the regulatory sequences (Fig. 1A-C). Seeking a better understanding of the actors regulating *Lmx1* expression in dorsal neural tube cells, we explore the *Ciona* single-cell transcriptomic atlas (Cao et al., 2019), focusing on the b-lineage nervous system from the initial gastrula to the mid gastrula stage, covering all stages of *Lmx1* expression and neural tube closure. After isolating these cells based on their expression of *Msx* and neural markers, we found that the b-lineage nervous system was divided into three states (Fig. S1A-D). Gene expression analysis indicates that one cell state corresponds to pLNPC-derived tail-tip cells, called rudder cells in this study, based on morphological similarities to the steering structure at the rear of a boat (Ishida and Satou, 2024). The two other groups are separated by *Cdx* expression, which might correspond to an anterior-posterior division of the b-derived dorsal neural tube domain (Imai et al., 2004). The *Cdx*^-^ and *Cdx*^+^ groups were named anterior and posterior dorsal neural tube cells, respectively (Fig. S1E-I). Notably, cells on the UMAP are arranged by stages. Early stage cells are present in a node branching into the three groups, while later stage cells progressively irradiate. The early and mid-tailbud stage were present at the periphery of these cell states (Fig S1J). In contrast to the anterior dorsal neural tube cells, *Lmx1* was sparsely expressed in posterior dorsal neural tube cells, reinforcing their assignment as posterior and the first to zipper (Fig. S1E). The cells were then ordered by pseudotime using the early stage cell or precursor cell cluster as the starting point (Fig. S1I). The single-cell trajectory branches into three transcriptional lineages corresponding to our three states (Fig. S1K). The transcription factors present in the posterior dorsal neural tube cells were then identified and ordered by pseudotime. Their expressions are separated into two waves: a first set that includes *Lmx1*, and a second set that includes *Msx*. These expression waves follow the documented expression patterns present in the dorsal neural tube cells (Ostlund-Sholars et al., 2025). In addition to Pax3/7 and Msx, the expression cascade contains multiple Zicl genes, transcription factors involved in early neural tube formation (Imai et al., 2002). The cells also express multiple Ets family transcription factors, which are involved in the Fgf/Erk signaling pathway, a major actor in *Ciona* nervous system development (Gainous et al., 2015; Hudson et al., 2003; Wagner and Levine, 2012). Finally, Smad effectors for both branches of the TGfβ superfamily signaling pathway are present in the cascade. Their binding motifs were then searched in the minimal enhancer sequence. It contains several motifs for Msx, Pax3/7, Zicl, Ets transcription factors, and Smads (Fig. 1A). To further analyze how these transcription factors could affect *Lmx1* expression, the binding motifs for each transcription factor were mutated. Loss of Msx or Ets binding motifs did not affect the reporter’s activity. The reporter loses all activity upon mutations of either Zicl or Smad binding motifs. On the contrary, reporter activity was increased, and even ectopic, when Pax3/7 binding motifs are mutated (Fig. 1B, 1D-E). Remarkably, *Zicl* transcription factors are present in the first wave of expression in the posterior cells (Fig S1L). Its knockdown using morpholino abrogates Lmx1 expression (Imai et al., 2006). *Smad2*/*3a* and *Smad1/5/9*, the nuclear effectors of the Nodal and BMP signaling pathway, respectively, share their binding motif sites. They are present in this first expression wave (Fig. S1L). It has not been shown that BMP can activate *Lmx1* expression in *Ciona,* but *Nodal* knockdown leads to *Lmx1* downregulation (Imai et al., 2006). This would represent a switch in the usage of the Tgfβ superfamily signaling pathway compared to vertebrates. Indeed, in vertebrates, the BMP signaling pathway activates *Lmx1* in the future roof of the neural tube (Chizhikov and Millen, 2004a, b). Zicl would then be a conserved regulator of *Lmx1* as it plays the same role in vertebrates (Elsen et al., 2008). The repressive effect of Pax3/7 was suggested by its complementary expression pattern to *Lmx1* (Fig. S1L). Its expression decreases as *Lmx1* expression increases and vice versa (Fig. S1L). Together, these data suggest that Zicl and Smad2/3 act as direct transcriptional activators while Pax3/7 represses *Lmx1* expression. These factors would account for the timely regulation of *Lmx1* in the dorsal neural tube.

**Fig. 1.**
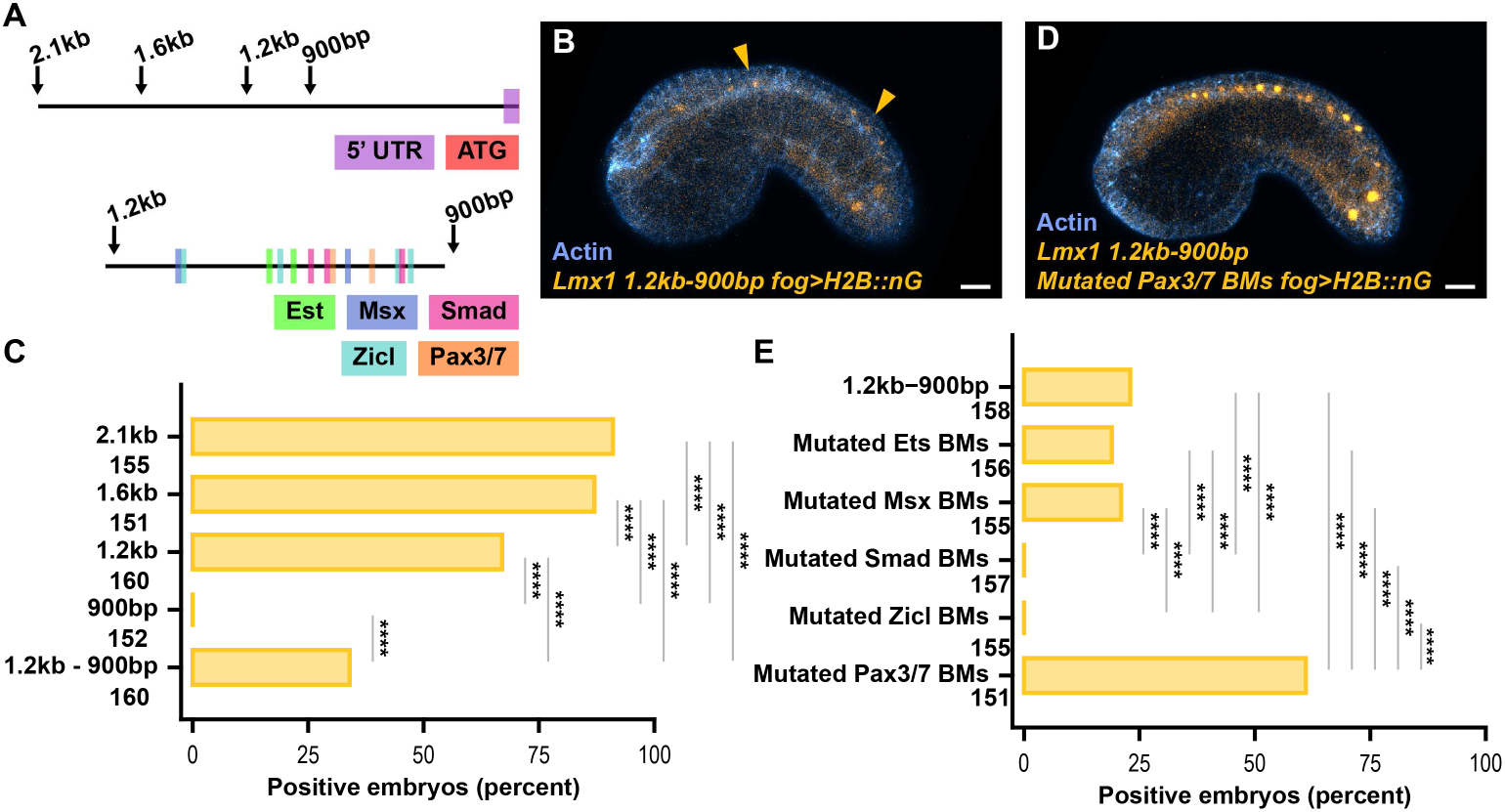
Transcriptional regulation of *Lmx1.* **(A)** Schematic of the tested regulatory sequences of the *Lmx1* locus (top) and of the minimal enhancer with the binding motif position color-coded by transcription factors (bottom). The length is from the translation start site. **(B)** Activity of the minimal enhancer of *Lmx1* (yellow, *Lmx1 1.2kb-900bp fog>H2B::nG*, nuclear reporter) in a representative mid-tailbud embryo stained for actin fibers (blue). Arrows point toward the dorsal neural tube. **(C)** Percentage of embryos expressing the regulatory sequences of different lengths in the dorsal neural tube cells (Chi square followed by Fisher test with Bonferroni correction: 2.1kb n = 155; 1.6kb n = 151; 1.2kb n = 160; 900bp n = 152; 1.2kb-900bp n = 160 total embryos (over 3 electroporations). **(D)** Representative image of a mid-tailbud embryo expressing nuclear reporter driven by the *Lmx1* minimal enhancer with mutated Pax3/7 binding motifs (*Lmx1 1.2kb-900bp Mutated Pax3/7 BMs fog>H2B::nG,* yellow) and counterstained for actin (blue). The pictures in B and in D have the same settings. **(E)** Percentage of embryos expressing the minimal enhancer with the different transcription factor binding motif mutations in the dorsal neural tube cells (Chi square followed by Fisher test with Bonferroni correction: 1.2kb-900bp n = 158; Ets mutated BMs, n = 156; Msx mutated BMs, n = 155; Smad mutated BMs, n = 157; Zicl mutated BMs, n = 155; Pax3/7 mutated BMs, n = 151 total embryos (over 3 electroporations)). **** p value <10^-4^. Scale bar: 20 µm.

### 2.2. Zippering defect in *Lmx1* knockdown

Misexpression of *Lmx1* is sufficient to disrupt neural tube closure (Ostlund-Sholars et al., 2025). To further investigate the role of *Lmx1* in neural tube closure and analyze if it is necessary for this process, we knock it down using morpholinos. Morpholinos were preferred over mutagenizing the *Lmx1* locus using Clustered regularly interspaced short palindromic repeats (CRISPR)/Cas9 due to *Lmx1* expression timing and the weak efficiency of Cas9 double stranded break at those stages, since CRISPR/Cas9 experiments in *Ciona* use F0 embryos (Gandhi et al., 2017; Gandhi et al., 2018; Stolfi et al., 2014). To visualize the neural plate border and the roof of the neural tube, a modified nuclear reporter for *Lmx1* resistant to the action of the morpholino due to mutations of its target site was injected along with the morpholino. Upon knockdown of *Lmx1*, the progression of the zippering fork in neurula embryos appears impaired and lagging compared to control injected embryos (Fig. 2A-B). To quantitatively assess this phenotype, the neuropore length was measured and normalized to the tail length to account for staging differences. It shows that the neural tube opening was slightly but significantly (p-value: 0.011) longer in *Lmx1* knockdown embryos than in control embryos (Fig. 2C). The *Lmx1* knockdown phenotype follows the pattern of its transcriptional regulators. Both *Nodal* signaling and *Pax3*/7 disruption cause neural tube closure defects (Kim et al., 2022; Mita and Fujiwara, 2007). Knockdown of *Zicl* is too disruptive to assess the effect on neural tube closure (Imai et al., 2006). However, its overexpression in the nervous system also leads to neural tube closure defect (Treen et al., 2023b). It is noteworthy that vertebrate orthologs of both *Pax3*/7 and *Zicl* have been implicated in tube closure (Epstein et al., 1991; Escuin et al., 2023; Inoue et al., 2004). Together, this suggests that vertebrate *Lmx1* orthologs might also contribute to neural tube morphogenesis, although multiple gene redundancies might mask this. In previous studies, no significant phenotype was observed in *Ciona Lmx1* knockdown embryos. It might be due to its subtlety. Indeed, the study reported expression patterns of key regulators for differentiation and might have overlooked slight defects in tube closure (Imai et al., 2006). Moreover, we cannot exclude the possibility that the phenotype is temporary and resorbs at a later stage. Nevertheless, this experiment shows that Lmx1 is involved in the proper progression of the zippering fork.

**Fig. 2.**
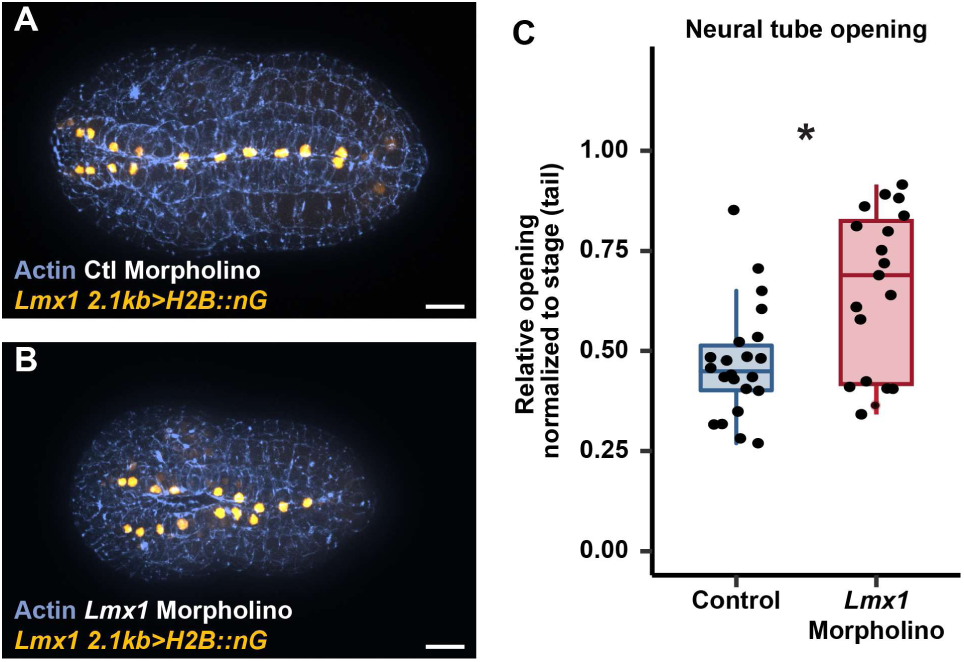
Neural tube defect upon downregulation of *Lmx1*. **(A**, **B)** Representative initial tailbud embryos injected with control or *Lmx1* morpholino and *Lmx1* nuclear reporter (*Lmx1>H2B::nG*, yellow) and counterstained for actin (blue). **(C)** Quantification of the neural tube opening normalized to stage as measured by the tail length. Dots are individual embryos. The box plot corresponds to the upper and lower quantiles, along with the median and the minimum and maximum values, excluding outliers. (Control, n = 25, *Lmx1* morpholino, n = 22 collected over 3 injection series, Wilcoxon rank-sum test, p value = 0.011). Scale bar 20µm.

### 2.3. Repressive action of Lmx1 during neural tube formation

In vertebrates, Lmx1 paralogs might act either as activators or repressors in a context-dependent manner (Chabrat et al., 2017; Chung et al., 2009; Doucet-Beaupre et al., 2016; Matsunaga et al., 2002; Yan et al., 2011). To assess the molecular mechanisms used by Lmx1 to regulate neural tube morphogenesis, a constitutively repressive Lmx1 variant (Lmx1::WRPW), a constitutively active form (Lmx1::VP64), or wildtype Lmx1 was overexpressed in dorsal neural cells using *Lmx1* regulatory sequences and compared to control embryos. The length of the neuropore at the mid-tailbud stage was again measured and normalized to the tail to account for stage differences. Upon overexpression of any *Lmx1* variant, the neural tube cannot close properly (Fig. 3A-E). However, there were drastic phenotypic differences. Indeed, as previously reported, overexpression of the constitutive form of Lmx1 keeps the tube mostly open, abrogating the zipper (Fig. 3A, 3D-E) (Ostlund-Sholars et al., 2025). In these cases, it seems that the b-muscle cells on the posterior tip of the tail and the rudder cells are transformed. These cells do not migrate ventrally. Instead, they form a U-shape rather than the normal V-shape fork. Their migration typically occurs shortly after the onset of the zipper movement (Hashimoto et al., 2015; Ishida and Satou, 2024). This defect possibly also prevents zippering progression. Overexpression of wild-type *Lmx1* in the dorsal cells causes zippering stalling at different points of the tube (Fig. 3A, 3C, 3E). Although significantly different, this phenotype was closer to that observed upon expression of the constitutively repressive *Lmx1* variant (Fig. 3A-B, 3E). Together, these data indicate that Lmx1 has both activating and repressive functions during neural tube morphogenesis, with possibly more targets repressed than activated.

**Fig. 3.**
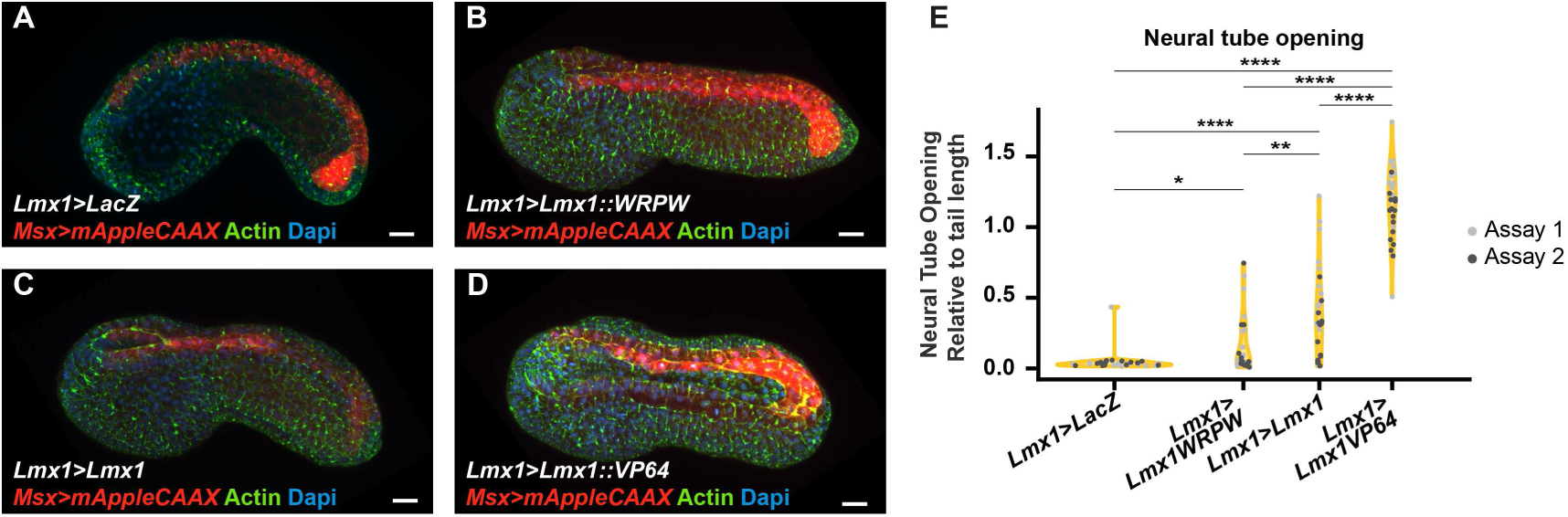
Effect of *Lmx1* variants on neural tube morphogenesis. **(A-D)**. Representative electroporated mid-tailbud embryos expressing either LacZ as controls (A, *Lmx1>LacZ*), a repressive motif (B, *Lmx1>Lmx1::WRPW*), wild-type Lmx1 (C, *Lmx1>Lmx1*), or Lmx1 fused to a constitutive activator domain (D, *Lmx1>Lmx1::VP64*) in the dorsal neural tube cells and a membrane reporter for Msx (*Msx>mAppleCAAX,* red). They were stained for cortical actin (green) and nuclei (Dapi, blue). (**E)** Quantification of the neural tube opening normalized to stage as measured by the tail length. Dots are individual embryos. (*Lmx1>LacZ:* n = 30 embryos; *Lmx1>Lmx1::WRPW*: n = 30 embryos; *Lmx1>Lmx1*: n = 30 embryos; *Lmx1>Lmx1::VP64*: n = 30 embryos, collected over two assays, Kruskal Wallis test followed by Wilcoxon rank sum test with Bonferroni correction: * p value < 0.05, ** p value < 0.01, **** p value < 0.0001. Scale bar 20µm.

### 2.4. Prevention of cellular intercalation by Lmx1

Given that our data points toward repressive functions of Lmx1 during neural tube morphogenesis, we extended our analysis of this variant. To gain insight into the mechanisms affected by the overexpression of this Lmx1 variant, initial tailbud embryos were analyzed. At this stage, all b-derived neural cells have zippered, and the a-derived dorsal neural cells will converge and fuse at the midline (Cole and Meinertzhagen, 2004; Hashimoto et al., 2015). At that moment, *Lmx1* is normally expressed anteriorly to the zippering fork and absent posteriorly (Ostlund-Sholars et al., 2025). Therefore, the Lmx1 repressive form is present in the posterior dorsal neural tube cells where endogenous Lmx1 is normally absent. Upon misexpression of the Lmx1 repressive form, embryos showed cell intercalation defects posterior to the zippering fork. Typically, dorsal cells begin intercalating as soon as the fork moves anteriorly, forming a one-cell dorsal row. This row is highlighted by the expression of the nuclear fluorescent Lmx1 reporter in dorsal cells that align along the midline. However, in embryos expressing the repressive *Lmx1* variant, two-cell rows were observed further from the zippering fork (Fig. 4). The absence of intercalation suggests that in the presence of the Lmx1 repressive form, genes normally involved in this process are not expressed. The stalling of the zipper might then be associated with the absence of intercalation. It is documented that issues in convergent extension, the alignment of lateral cells toward the midline, associated with planar cell polarity mutations, lead to neural tube closure defects in vertebrates (De Marco et al., 2014; Galea et al., 2018; Kibar et al., 2001; Wallingford and Harland, 2002). However, no neural tube closure defect has been reported in planar cell polarity mutants in *Ciona.* This discrepancy might be associated with the absence of neural fold elevation in Ciona prior to the closure (Hashimoto et al., 2015). Alternatively, it might have been overlooked due to the drastic notochord malformation in these species (Jiang et al., 2005; Shi et al., 2009). Mutations affecting positively or negatively the planar cell polarity pathway usually led to similar phenotypes. This phenomenon might explain why Lmx1 knockdown and the overexpression of its repressive form cause neural tube closure defects. Moreover, in *Ciona*, the driving forces for tail elongation are mostly associated with the intercalation of notochord cells (Sehring et al., 2014). The rest of the tail continues to elongate, leading to an imbalance in diverse tissue forces and overriding the normal mechanism of tail bending (Kogure et al., 2022; Kogure et al., 2026; Lu et al., 2020). It causes the tail to stay straight or to bend upward, as observed in *Lmx1* misexpression experiments (Fig. 3A-D). To conclude, *Lmx1* disruption triggers a morphogenetic imbalance during neural tube formation.

**Fig. 4.**
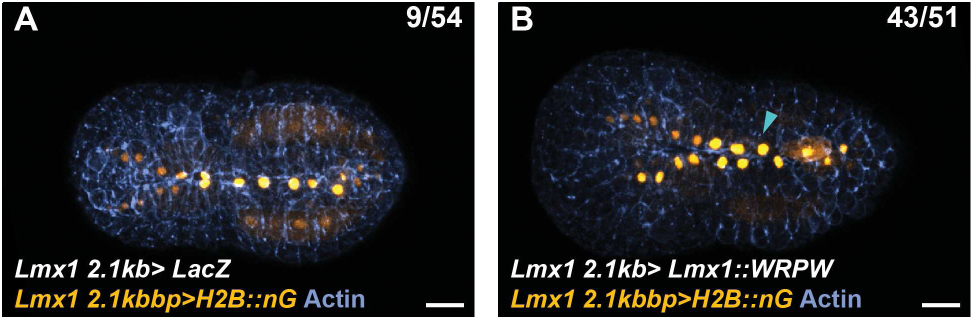
Intercalation defect in Lmx1 repressive variant. **(A,B)** Representative images of initial tailbud embryos expressing a nuclear reporter (*Lmx1>H2B::nG,* yellow) and *LacZ* as control (F, *Lmx1>LacZ*) or repressive *Lmx1* (G, *Lmx1>Lmx1::WRPW*) in the dorsal neural tube cells using the *Lmx1* driver and stained for cortical actin (blue). Cyan arrowheads point to not intercalating cells in G. The number of embryos in the upper left corner indicates the number of intercalation defects (cyan arrowheads) over the total number of embryos (pooled over 3 electroporations). Scale bar 20µm.

### 2.5. *Laminin* regulation in embryos expressing constitutively repressive Lmx1 variants

Lastly, we investigated potential downstream targets of Lmx1 associated with morphogenesis. Using our single-cell trajectory of posterior dorsal neural tube cells, we found that these cells dynamically express the extracellular matrix component *Laminin* (Table S1). Laminin is known to play a role in notochord cell intercalation in *Ciona*. Spontaneous gene mutation causes the notochord cells to fail to intercalate (Veeman et al., 2008). Based on the single-cell trajectory, its expression level increases over time, opposite to *Lmx1* expression (Fig. 5A, S1L). To test the hypothesis that *Laminin* expression is regulated by *Lmx1,* we perform in situ hybridization chain reaction (HCR) for *Laminin* mRNA on electroporated embryos expressing *LacZ*, as control, or repressive *Lmx1* in the dorsal neural tube cells. The embryos were co-electroporated with a nuclear *Lmx1* reporter to distinguish transgenic nuclei from those that did not receive the plasmids. Our results showed that in the presence of the repressive *Lmx*1 variant, nascent Laminin mRNA expression decreased. It was observed both between controls and *Lmx1* repressive embryos and within mosaic embryos expressing the repressive *Lmx1* form (Fig. 5B-E). Indeed, nuclei without misexpression have a significantly higher level of nascent *laminin* transcript (Fig. 5F). These data indicate that *Lmx1* is able to regulate Laminin. It also suggests that *Lmx1* might prevent the early expression onset of genes involved in the latter step of neural tube morphogenesis, such as *Laminin*. Early expression of later genes might impede zipper progression, explaining the *Lmx1* morpholino phenotype. Therefore, Lmx1 would be required to actively repress these genes and should be cleared to allow the later step to proceed.

**Fig. 5.**
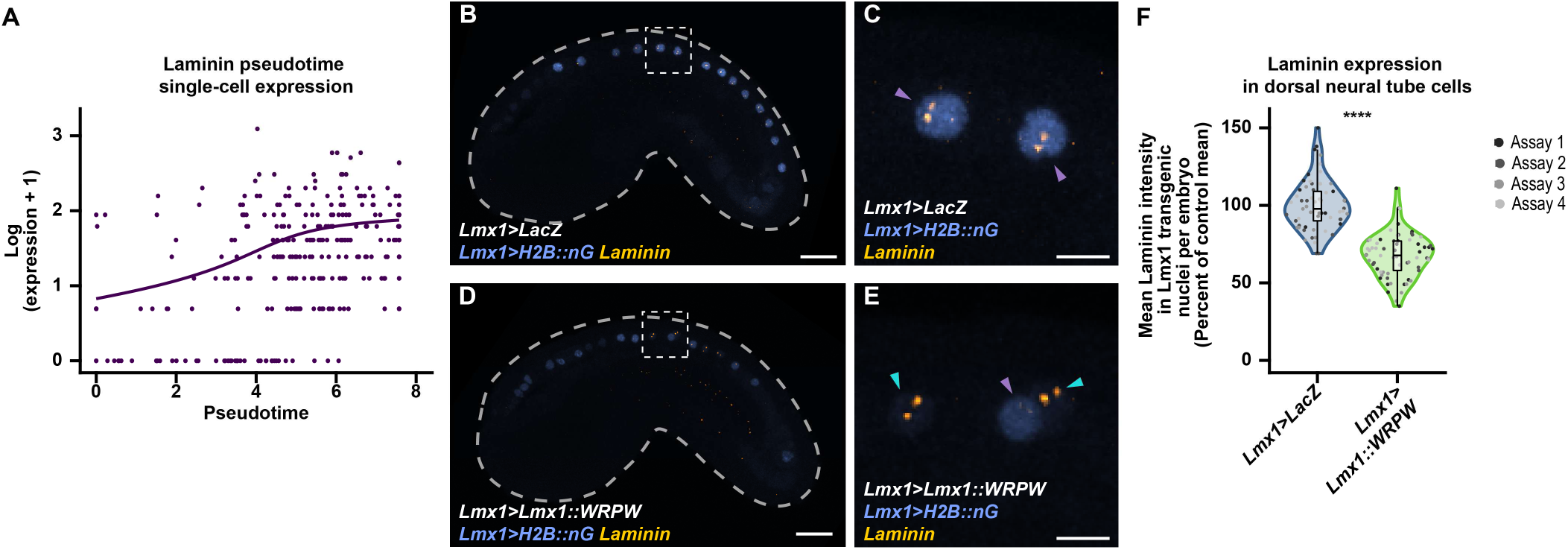
*Laminin* regulation by constitutively repressive Lmx1. **(A)** Single cell expression of *Laminin* in the posterior dorsal neural tube cells along pseudotime (n = 398 cells). (**B-E)** Representative images of HCR for *Laminin* mRNA (yellow) on electroporated mid-tailbud embryos expressing LacZ as controls (B, C, *Lmx1>LacZ*) or repressive *Lmx1* (D, E, *Lmx1>Lmx1::WRPW*) and a nuclear *Lmx1* reporter (*Lmx1>H2B::nG*, blue) in the dorsal neural tube cells. C and E are magnifications of the dashed area in B and D, respectively. Gray dashed lines outline the embryo. Purple arrowheads point to transgenic *Lmx1* nuclei and cyan to non-transgenic nuclei. **(F)** Violin plot with box plot of the normalized laminin fluorescence intensity in transgenic *Lmx1* nuclei per embryo, shown in B to D. Dots represent individual embryos, color-coded per assay. Box plots show the minimum, first quartile, median, third quartile, and maximum. (Control: n = 61, *Lmx1>Lmx1::WRPW:* n = 65, collected over 4 electroporations, Wilcoxon rank-sum test, p value <10^-4^). Scale bar 20µm (B, D) and 5µm (C, E).

**Fig. 5.**
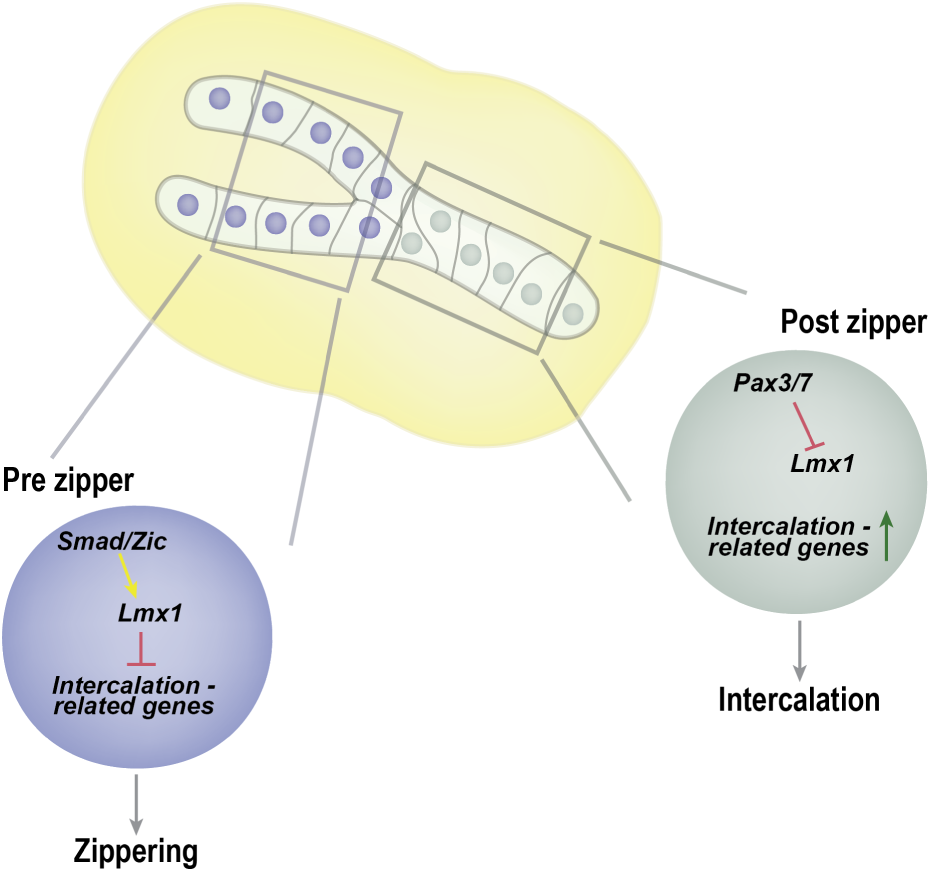
Proposed Model of Lmx1 role as temporal gatekeeper during neural tube morphogenesis. In *Ciona* neurulating embryos, Lmx1 is expressed in the dorsal neural cells in front of the zippering fork. In these cells, *Lmx1* is activated by Smad2/3 and Zicl factors. It might repress intercalation genes. Posterior to the zippering fork, *Lmx1* is repressed by Pax3/7. Intercalation genes are therefore expressed. This mechanism enables the timely coordination of morphogenesis.

## 3. Conclusion

In this study, we uncovered how *Lmx1* is regulated in the dorsal neural tube cells of *Ciona* embryos. We found that Zicl and Smad factors directly activate its expression, while Pax3/7 might repress it (Fig. 5). This tight regulation allows stringent control of *Lmx1* functions. We further highlight the involvement of *Lmx1* in neural tube closure using knockdown and misexpression experiments. In this process, Lmx1 might act as a repressor in the future dorsal cells that have not yet zipped. Its presence might prevent the expression of genes involved in later steps of development, such as *Laminin*, allowing for a smooth zippering transition. Therefore, it suggests *Lmx1* can coordinate developmental timing, acting as a morphogenetic gatekeeper (Fig. 5). Future experiments will elucidate the extent of Lmx1’s gatekeeping role, how many target genes it represses, how it coordinates both cell cycle and morphogenesis, and whether this function might be conserved with vertebrates.

## 4. Methods

### 4.1. Ciona handling

Marinus and M-Rep collected adult *Ciona intestinalis* type A (Pacific species, also called *Ciona robusta)* along the shorelines of Long Beach (CA, United States) and San Diego (CA, United States), respectively. They were shipped overnight and kept in artificial seawater under constant illumination at 18°C until we collected their gametes. Eggs were dechorionated with a protease solution (Protease from Streptomyces griseus, Sigma). These procedures were performed using the protocol described by Christian and colleagues (Christiaen et al., 2009b).

### 4.2. Electroporation

Fertilized dechorionated eggs were mixed with plasmid DNA and electroporated, according to published instructions (Christiaen et al., 2009a; Corbo et al., 1997). Plasmid DNA mixes for electroporation are given in Table S2. Once electroporated, the embryos were reared in filtered artificial seawater between 18 and 22°C until they reached the desired stage. They were fixed in 4% formaldehyde, 0.1 M Mops (pH 7.4), 0.5 M NaCl, 1 mM EGTA, 2 mM MgSO4, and 0.05% Tween 20 for 30 min at room temperature. Once fixed, embryos were washed several times in PBST (phosphate-buffered saline (PBS) with 0.01% Tween 20). They might be stained with phalloidin conjugated to a fluorochrome (ActinGreen, Thermofisher; ActinRed, Thermofisher; Phalloidin CF^®^660R, Thomas Scientific), the nuclear dye 4′,6-diamidino-2-phenylindole (Dapi, Thermofisher), or both. Embryos were then mounted on slides using Fluosave mounting medium (Sigma). Electroporation assays were performed at least twice using different egg batches.

### 4.3. Microinjection

Dechorionated eggs were injected with a mixture of linearized plasmids, 1mg/ml Dextran, Alexa Fluor™ 647, 10,000 MW (D22914, ThermoFisher), and control or *Lmx1* morpholinos using pulled glass needles mounted on a pressure injector (World Precision Instruments). For the injection, eggs were placed in agarose-cast microwells (Gregory and Veeman, 2013). Morpholinos (*Lmx1*: GAAGAACGCAGCATTTAACACTAGA) and random control oligo 25-N were produced by Gene Tools, LLC (Philomath, OR). Mixes are described in Table S2. Eggs were fertilized, and embryos were then grown at 22°C until control embryos reached the initial tailbud stage. They were fixed using the same procedure as for electroporation and stained with phalloidin CF^®^660R and Dapi. Microinjections were performed in triplicate with different egg batches.

### 4.4. Cloning

Gene names, their KH and KY gene model identifiers and common synonyms mentioned in this study are given in Table S3. The 2.1kb *Lmx1* regulatory sequences controlling the expression of histone 2B (H2B) fused to either the fluorescent protein neongreen (*H2B::nG*) or mApple (*H2B::mAp*) in the pCESA expression vector were previously described (Lemaire et al., 2021; Shaner et al., 2013; Shaner et al., 2008; Todorov et al., 2024). 5’ truncated regulatory sequences were obtained by PCR amplifying different fragments of the 2.1kb *Lmx1* regulatory sequences (PCR primers in Table S4), digesting using AscI and NotI restriction enzymes (New England Biolab, NEB), and ligating them with T4 DNA ligase (Promega) into the same expression vector. The expression vector containing the *Lmx1* minimal enhancer fragment was obtained by recombination using NEBuilder® HiFi DNA Assembly Cloning Kit (NEB). Fragment of the regulatory sequence was PCR-amplified from the *Lmx1 1.2kb>H2B::mApple* (fragment #1, primers in Table S4) and recombined to a backbone containing the fog minimal promoter upstream of *H2B::mAp*, which was PCR-amplified to obtain the correct overhangs (fragment #2, primers in Table S4) (Cao et al., 2019; Rothbacher et al., 2007). Mutagenesis of binding motifs contained in the *Lmx1* minimal enhancer was performed by rounds of recombination of PCR-amplified fragments from the original plasmid using primers harboring the mutated nucleotides (recombination series and primers in Table S4). Minimal enhancer sequences are provided in the Supplementary file S1. *Lmx1* reporter resistant to the morpholino (by modifying the morpholino complementary sequence to atgTtAAgAgcggc) or containing the full complementary sequence were obtained by editing the morpholino binding site using recombination of PCR amplified fragment with the appropriate overhang (primers in Table S4).

The regulatory sequence of *Msx* was published by Abitua and colleagues (Abitua et al., 2012). It was cloned in the same expression vector upstream of mApple targeted to the membrane by the palmytoylation motif CAAX (*mAp-CAAX*) using ligation after restriction enzyme digestion with NotI and AscI (NEB) (Lemaire et al., 2021).

The Lmx1 misexpression pCESA plasmid *Lmx1>Lmx1::VP64* and the pCESA *LacZ* expression plasmid controlled by Lmx1 regulatory sequences (*Lmx1>LacZ*) were described by Ostlund-Sholar and colleagues (Fujiwara et al., 1998; Ostlund-Sholars et al., 2025). The *Lmx1* regulatory sequence was digested with NotI and AscI and ligated into a pCESA expression plasmid upstream of the coding sequence of *Lmx1* digested with the same restriction enzymes (Ostlund-Sholars et al., 2025). The repressor motif WRPW (Wagner et al., 2014) was inserted downstream of the *Lmx1* open reading frame by PCR amplifying *Lmx1>Lmx1* expression plasmid with primers containing the motif (Table S4), digesting with DpnI restriction enzyme to remove the template (NEB), and recombining the fragment.

### 4.5. HCR

Electroporated mid-tailbud embryos were fixed overnight at 4°C with a solution composed of 100 mM HEPES, 500 mM NaCl, 1.75 mM MgSO4, 2 mMethylene glycol bis (succinimidyl succinate) (Thermo Fisher Scientific, 21565), and 1% formaldehyde (Thermo Fisher Scientific, 28908) (Treen et al., 2023a). The next day, the samples were washed several times with PBST, dehydrated in 50% ethanol, then 80% ethanol, and stored at −20°C. After rehydration in PBST, embryos were hybridized with Laminin oligoprobes purchased from Molecular Instruments following their v3.0 RNAFISH HCR protocol for sea urchin (Choi et al., 2016). Briefly, after blocking, the embryos were incubated overnight at 37°C in the hybridization buffer with the probe. The embryos were then washed to remove excess probes. Probe detection and signal amplification were done with the fluorochrome-conjugated hairpin amplification, B2-Alexa 546 (Molecular Instruments), for three hours in the dark at room temperature. After the final wash, the embryos were mounted on glass slides with RapiClear 1.49 (Sunjin Lab).

### 4.6. Image acquisition and analysis

Zeiss 880 confocal microscope (Zen Black v.2.3SP1 acquisition software), Zeiss Apotome (ZEN 3.10 acquisition software), and Leica Sp8 scanning confocal microscope (LAS X version v3.5.7.23225 acquisition software) were used to image fixed embryos. Raw images produced with the Zeiss apotome were processed in Zen Blue to generate virtual optical images. Most analyses and maximum intensity around the dorsal cell of the neural tube were performed in ImageJ/Fiji (version 2.14.0/1.54j) (Schindelin et al., 2012). The Pearson Chi square followed by the Fisher Exact Test with Bonferroni correction was used to test differences between embryos expressing the various Lmx1 reporters. Measurement of neural tube opening was performed in FluoRender (2.30/2.34). Neural tube opening length was normalized to tail length to acknowledge stage differences. Statistical differences between Lmx1 morpholino and control embryos were tested using the nonparametric Wilcoxon rank-sum test. Differences between the Lmx1 variant misexpression conditions were assessed with pairwise Wilcoxon rank-sum tests with Bonferroni correction as post hoc tests of the Kruskal-Wallis statistical test. To analyze *laminin* mRNA expression between controls and embryos expressing *Lmx1::WRPW*, the nuclei expressing the *H2B::nG* transgenes were segmented. The fluorescence intensity of the laminin signal was then measured within each segmented nucleus as readout of nascent transcription. Segmentation and measurement were performed with a custom script available on GitHub. The intensity was then averaged for each embryo to account for electroporation mosaicism. These fluorescence intensity averages were then compared between control and *Lmx1::WRPW* embryos. The difference was then statistically tested using the nonparametric Wilcoxon rank-sum test. Graphs and the statistical analysis were done in R (v4.5.1) using the ggplot2 package (v4.0.2) and the stat package (R Core Team, 2022; Wickham, 2009).

### 4.7. Single cell analysis

Reads from the Cao et al study from the initial gastrula to the mid tailbud stage available on SRA (dataset: SRX5827302, SRX5827303, SRX5827304, SRX5827305, SRX5827306, SRX5827307, SRX5827308, SRX5827309, SRX5827310, SRX5827311, SRX5827312, SRX5827313, SRX5827314, SRX5827315; project: PRJNA542748) were remapped on the KY21 gene model using Cell Ranger (v 7.0.1). Single-cell RNA analysis was then performed in R (v4.5.1) with Seurat v5.4.0 (Stuart et al., 2019). Single-cell expression matrices were filtered. Cells with more than 1000 genes, between 0.08% and 20% mitochondrial gene content, and less than UMI per dataset of average + 5 standard deviations were selected for further analysis. The read counts were then log-normalized and integrated. The cells were then clustered using the Leiden clustering algorithm to identify the nervous system, which was then subsetted (Traag et al., 2019). The b-lineage was then subsetted based on neural Msx expression and reclustered. Additional epithelial cells were then removed, based on their Uxs1 expression. After each subsetting, cells were reintegrated and reprojected using UMAP. Once the posterior dorsal neural tube was identified, Slingshot (v2.18.0) was used to perform pseudotime analysis on the final UMAP projection (Street et al., 2018). Dynamics gene expression analysis was performed using tradeSeq (v1.24.0) (Van den Berge et al., 2020). The expression level of transcription factors with more than 4 counts in at least 12 cells was then plotted along pseudotime using complexHeatmap (v2.26.1) (Gu et al., 2016).

### 4.8. Binding motif analysis

Position Weight Matrices (PWM) for the binding motifs for Pax3/7, Msx, Ets1/2.b, Erf.b, Elk, Smad1/5/9, Smad2/3.a, Zicl1, Zicl3, Zicl4, Zicl5, Zicl6, and Macho were obtained from Systemic Evolution of Ligands by Exponential Enrichment (SELEX) assays published by Nitta and colleagues (Nitta et al., 2019). If the binding motif for the given transcription factor was not identified in this SELEX study, the PMW for its most similar human transcription factor was retrieved from JASPAR database (Ovek Baydar et al., 2026). PMW are provided in the supplementary file S2. This list was then used to scan the *Lmx1* minimal enhancer sequence for binding sites using motif enrichment analysis FIMO (Find Individual Motif Occurrences) (Grant et al., 2011). Identified motifs are provided in Table S5. The motifs with the 2 to 4 lowest p-value for each transcription factor family were then mutated in the minimal enhancer.

## CRediT authorship contribution statement

Josefina Perez: Formal analysis, Investigation, Visualization, Writing – original draft. Michael S. Levine: Conceptualization, Funding acquisition, Supervision, Writing – review and editing. Laurence. A. Lemaire: Conceptualization, Formal analysis, Funding acquisition, Investigation, Methodology, Supervision, Visualization, Writing – original draft.

## Acknowledgements

The authors would like to thank the members of the Lemaire Lab and of the Levine Lab, especially Dr. Anrea Mariossi and Dr. Nicholas Treen. They also extend their acknowledgments to Dr. Mohini Sengupta from Saint Louis University for helpful discussion. A grant (NS076542) from the National Institute of Neurological Disorders and Stroke of the National Institutes of Health (NIH) to M.S.L supported the initial phase of the project presented in this manuscript. This research was also supported by the Office of Vice President for Research and the Provost Office of Saint Louis University (RI award 01978 and grant 001654) to L.A.L.

## Data availability

The reads single-cell RNA datasets were already published (Cao et al., 2019). They are deposited in Sequence Read Archive (SRA) under the accession number: SRX5827302, SRX5827303, SRX5827304, SRX5827305, SRX5827306, SRX5827307, SRX5827308, SRX5827309, SRX5827310, SRX5827311, SRX5827312, SRX5827313, SRX5827314, SRX5827315 (project number: PRJNA542748). R scripts for single-cell analysis and custom Fiji scripts are available on GitHub (https://github.com/LemaireLabAtSLU/Ciona-Lmx1)

---

**Fig. S1.**
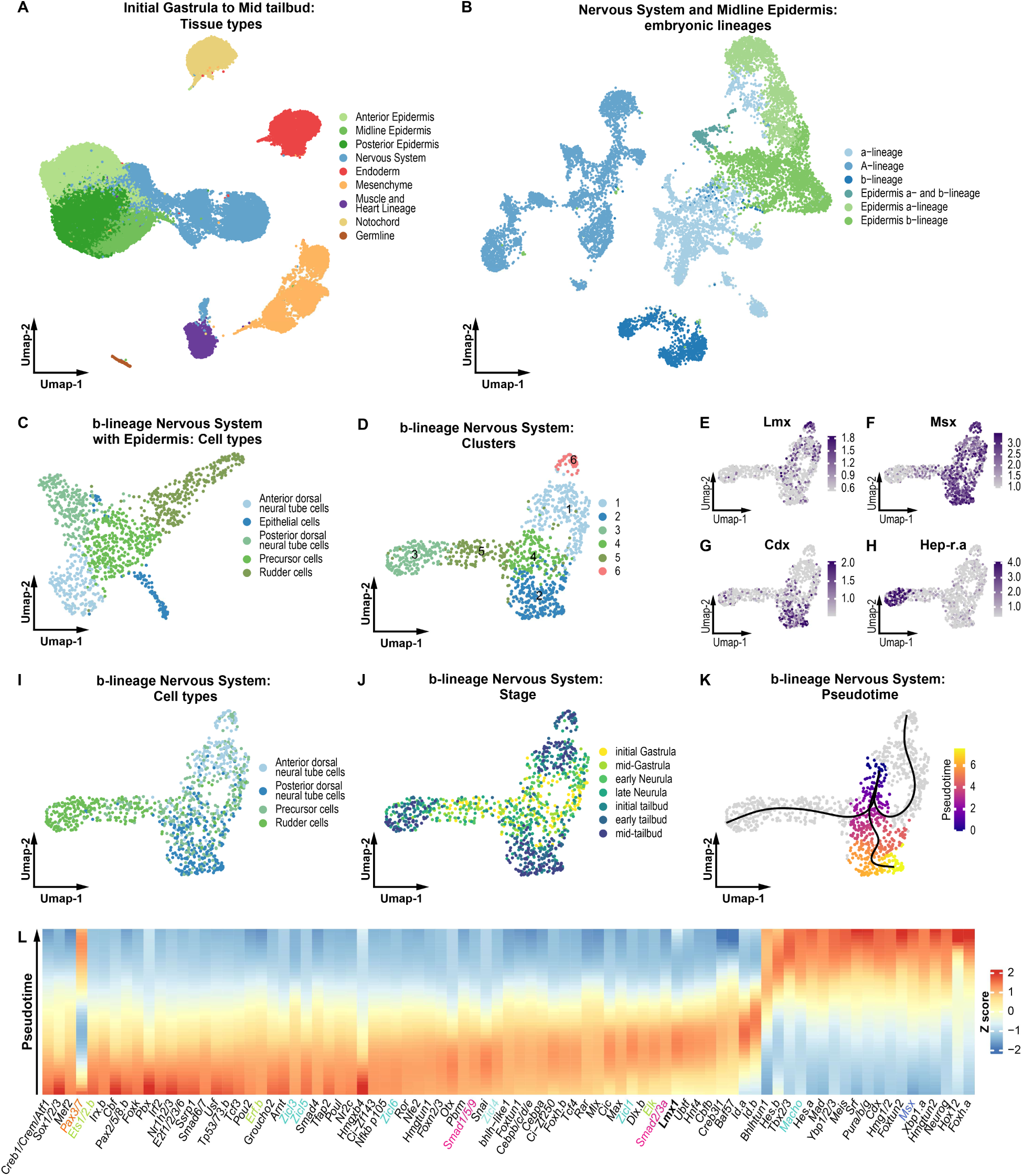
Single cells transcriptomic analysis of the b-nervous system. **A.** Uniform Manifold Approximation and Projection (UMAP) of the *Ciona* single cells from initial gastrula to mid tailbud stage, totaling 7 stages from the Cao et al., 2019 datasets. Cells were color-coded by tissues, with the epidermis divided into anterior, midline, and posterior (n = 43457 cells). **B.** UMAP of the nervous system and midline epidermal cells shown in A. and color-coded based on lineages and tissue type (n = 11917 cells). **C.** UMAP of the b-lineage nervous system shown in B. and color-coded by cell types (n = 1107 cells). **D.** UMAP of the cells shown in C, except the epidermal cells, and cluster color-coded (n = 1021 cells). **E-H.** Expression level of *Lmx1* (E), *Msx* (F), *Cdx* (G), and *Hep-r.a* (H). on the UMAP presented in D. **I-J**. UMAP of the cells shown in D. and color-coded by cell types (I) or stage (J) (n = 1021 cells). **K.** UMAP of the cells shown in D harboring the pseudotime color code of the posterior dorsal neural tube cells corresponding to clusters 4 and 2 in J. Black lines correspond to the single-cell trajectory of the three transcriptomic lineages identified for the b-nervous system (n = 1021 cells). **L.** Heatmap showing the expression level of the transcription factors present in the posterior dorsal neural tube cells and aligned by pseudotime (n = 398 cells). Further studied transcription factors are colored the same as Fig. 1.

